# Validation of a novel associative transcriptomics pipeline in *Brassica oleracea:* Identifying candidates for vernalisation response

**DOI:** 10.1101/2020.11.27.400986

**Authors:** Shannon Woodhouse, Zhesi He, Hugh Woolfenden, Burkhard Steuernagel, Wilfried Haerty, Ian Bancroft, Judith A. Irwin, Richard J. Morris, Rachel Wells

**Affiliations:** Department of Crop Genetics, John Innes Centre, NR47UH Norwich, UK; Department of Biology, University of York, Heslington, YO105DD York, UK; Computational & Systems Biology, John Innes Centre, NR47UH Norwich, UK; Earlham Institute, NR47UH Norwich, UK; School of Biological Sciences, University of East Anglia, NR47TJ Norwich, UK

**Keywords:** GWAS, Population Structure, Associative Transcriptomics, Flowering, Vernalisation, *Brassica oleracea*

## Abstract

Associative transcriptomics has been used extensively in *Brassica napus* to enable the rapid identification of markers correlated with traits of interest. However, within the important vegetable crop species, *Brassica oleracea*, the use of associative transcriptomics has been limited due to a lack of fixed genetic resources and the difficulties in generating material due to self-incompatibility. Within Brassica vegetables, the harvestable product can be vegetative or floral tissues and therefore synchronisation of the floral transition is an important goal for growers and breeders. Vernalisation is known to be a key determinant of the floral transition, yet how different vernalisation treatments influence flowering in *B. oleracea* is not well understood.

Here, we present results from phenotyping a diverse set of 69 *B. oleracea* accessions for heading and flowering traits under different environmental conditions. We developed a new associative transcriptomics pipeline, and inferred and validated a population structure, for the phenotyped accessions. A genome-wide association study identified miR172D as a candidate for the vernalisation response. Gene expression marker association identified variation in expression of *BoFLC*.C2 as a further candidate for vernalisation response.

This study provides insights into the genetic basis of vernalisation response in *B. oleracea* through associative transcriptomics and confirms its characterisation as a complex G x E trait. Candidate leads were identified in miR172D and *BoFLC*.*C2*. These results could facilitate marker-based breeding efforts to produce *B. oleracea* lines with more synchronous heading dates, potentially leading to improved yields.

## Introduction

Ensuring synchronous transiting from the vegetative to the reproductive phase is important for maximising the harvestable produce from brassica vegetables. Many cultivated brassica vegetables arose from their native wild form *B. oleracea* var. *oleracea*^1^. Wild cabbage, *B. oleracea* L., is a cruciferous perennial growing naturally along the coastlines of Western Europe. From this single species, selective breeding efforts have enabled the production of the numerous subspecies we see today. The specialization of a variety of plant organs has given rise to the large diversity seen within the species. Various parts of brassicas are harvested, including leaves (e.g. leafy-kale and cabbage), stems (e.g. kohl-rabi), and inflorescences (broccoli and cauliflower). For all subspecies, the shift from the vegetative to reproductive phase is important and being able to genetically manipulate this transition will aid the development and production of synchronous brassica vegetables.

Determining how both environmental and genotypic variation affect flowering time is important for unravelling the mechanisms behind this transition. For many *B. oleracea* varieties, a period of cold exposure, known as vernalisation, is required for the vegetative-to-floral transition to take place. This requirement for vernalisation, or lack thereof, determines whether the plant is a winter annual, perennial or biennial or whether it is rapid-cycling or a summer annual^2^. As a consequence, the response of the plant to vernalisation provides quantifiable variation that has been exploited by breeders to develop varieties with more synchronous heading. Such variation will be key for future breeding in the face of a changing climate.

Genome-wide association studies (GWAS) are an effective means of identifying candidate genes for target traits from panels of genetically diverse lines^3^. GWAS has been used successfully in numerous plant species including Arabidopsis, maize, rice and Brassica^4–7^. However, its application is reliant on genomic resources which are not always available for complex polyploid crops. Associative transcriptomics uses sequenced transcripts aligned to a reference to identify and score molecular markers that correlate with trait data. These molecular markers represent variation in gene sequences and expression levels. Associative transcriptomics is a robust method for identifying significant associations and is being used increasingly to identify molecular markers linked to trait-controlling loci in crops^8–12^.

An important factor to account for in association studies is the genetic linkage between loci. If the frequency of association between the different alleles of a locus is higher or lower than what would be expected if the loci were independent and randomly assorted, then the loci are said to be in linkage disequilibrium (LD)^13^. LD will vary across the genome and across chromosomes and it is important to account for this in GWAS analyses. This variation in LD is due to many factors, including selection, mutation rate and genetic drift. Strong selection or admixture within a population will increase LD. Accounting for the correct population structure reduces the risk of detecting spurious associations within GWAS analyses. The population structure can be determined from unlinked markers^14^.

Here, we develop and validate an associative transcriptomics pipeline for *B. oleracea*. A specific population structure consisting of unlinked markers was generated using SNP data from 69 lines of genetically fixed *B. oleracea* from the Diversity Fixed Foundation Set^15^. The pipeline was successfully used for the identification of candidate leads involved in vernalisation response, identifying a strong candidate in miR172D.

## Results

### Exposure to different environmental conditions identifies vernalisation requirements across the phenotyped accessions

We selected a subset of 69 *B. oleracea* lines, diverse in both eco-geographic origin and crop type, from the *B. oleracea* Diversity Fixed Foundation Set^16^. We used these accessions to evaluate the importance of vernalisation parameters by quantifying flowering time under different conditions (vernalisation start, duration and temperature). Two key developmental stages were monitored: ‘days to buds visible’ (DTB) and ‘days to first flower’ (DTF). The variation in flowering time across the different treatments and between the different lines is shown in Fig. 1. The different vernalisation start times demonstrate that exposure to the longer, ten-week pre-vernalisation growth period typically results in earlier flowering, compared to the shorter, six-week growth period. The mean DTB for the six-week growth period was 21.0 days (SD = 51.6), compared to 5.8 days (SD = 49.9) for the ten-week period (Wilcoxon Test, W = 17958, P = 0.004). Similarly, we found a significant difference in the time taken to reach DTF between the two treatment groups, with a mean of 57.9 days (SD = 55.5) following the six-week pre-growth period, in comparison to 35.9 days (SD = 53.1) following the ten-week pre-growth period (Wilcoxon Test, W = 17471, P = 2.96e-05).

**Figure 1:**
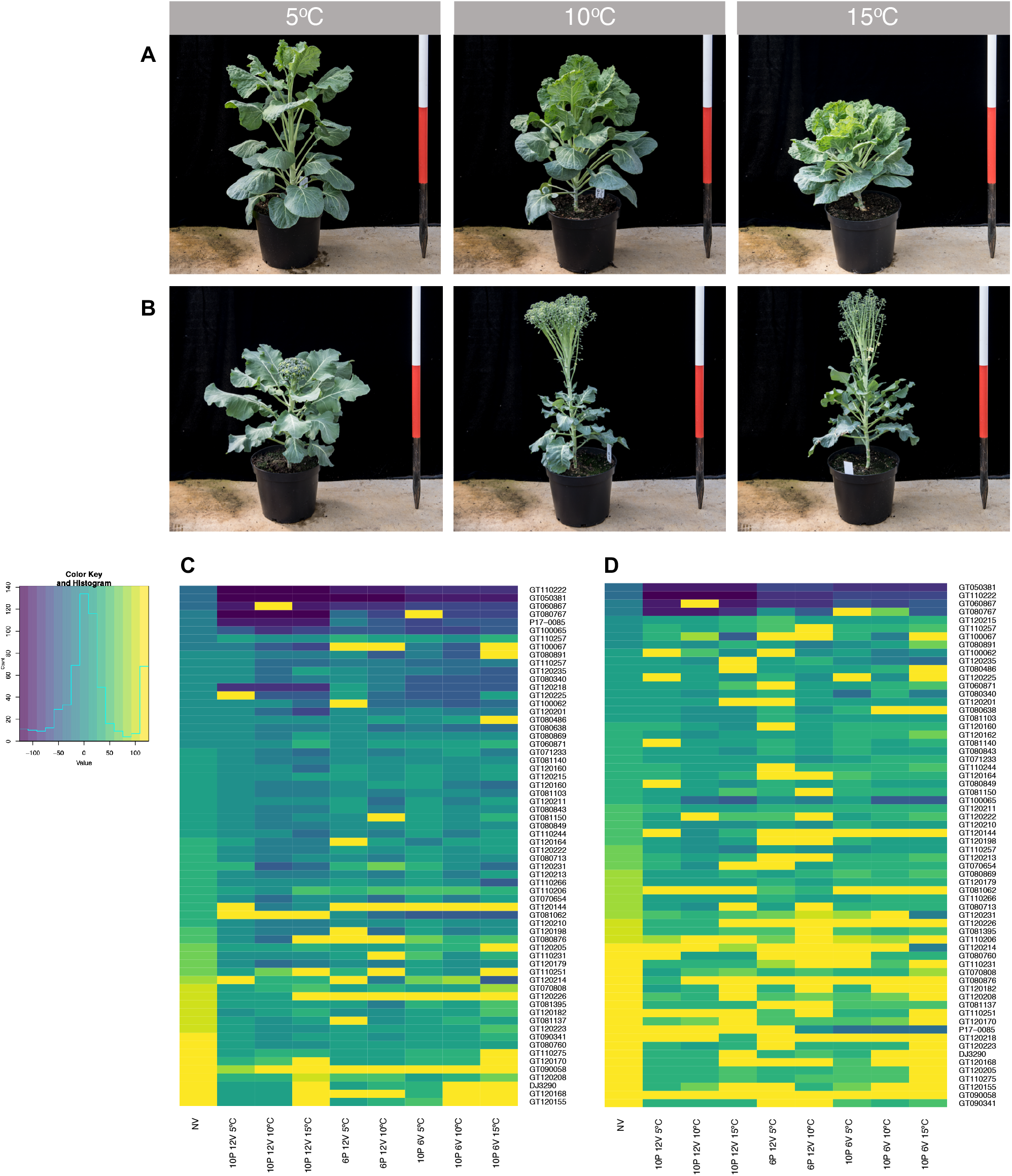
Flowering time traits exhibit a varied response to different environmental conditions within the population. A) Phenotypic response to different vernalisation temperatures in the Brussels Sprout Cavolo Di Bruxelles Precoce (GT120168). B) Phenotypic response to different vernalisation temperatures in the Broccoli Mar DH (GT110244). C) DTB per treatment, per line. D) DTF per treatment, per line.

Changes in vernalisation duration led to a significant difference in DTB, but not in DTF. Following the six-week vernalisation treatment, the mean DTB was 9.5 days (SD = 44.5) compared to 5.8 days (SD = 46.8) after exposure to twelve-weeks of vernalisation (Wilcoxon Test, W = 19532, P = 0.002). This difference was coupled with more synchronous heading between lines following the twelve-week vernalisation period. The impact of vernalisation duration on DTB varied across the population, reflecting the numerous factors that can affect DTB depending on crop type, such as stem elongation and developmental arrest.

Of the three parameters we investigated, vernalisation temperature resulted in the most pronounced phenotypic differences. The 5 °C vernalisation resulted in the largest DTB (slowest overall bud development), whereas the 10 °C treatment resulted in the largest DTF. The distribution between heading dates was distinctly different between the temperatures. Higher vernalisation temperatures resulted in larger the variation in DTB and DTF. The more synchronous heading and flowering for the 5 °C treatment suggests that this temperature was able to saturate the vernalisation requirement for a large proportion of the lines. After exposure to the warmer temperatures, the variation in DTB and DTF were greatly increased (Supplemental Fig. 1), indicating that the cooler vernalisation temperature aided faster transitioning in some lines, but delayed the development of others. This is consistent with differences in *B. oleracea* crop types, for example Brussels Sprouts are known to have a strong vernalisation requirement, whereas Summer Cauliflower have been bred to produce curd rapidly without the need for cold exposure^17,18^.

The effect of vernalisation temperature on the floral transition is demonstrated clearly between the Broccoli Mar DH and the Brussel Sprout Cavolo Di Bruxelles Precoce (Fig.1A), with polar responses to vernalisation temperature. Mar DH transitioned fastest under the 15 °C vernalisation treatment, whereas Cavolo Di Bruxelles Precoce transitioned faster under the 5 °C vernalisation treatment. Faster transitions at higher vernalisation temperatures as in the case of Mar DH, however, can lead to undesirable phenotypes from a grower’s perspective (Fig. 1B).

### Unlinked markers are required to generate a representative population structure

GWAS requires trait, SNP and population data. The correct population structure is important for ensuring that associations are with the trait of interest rather than identified on account of relatedness within the population, in particular for panels of only one species. To generate a representative population structure, it is necessary to ensure the SNPs used are unlinked^14^. However, different criteria have been used to select these SNPs^6,19–21^. To evaluate the impact of SNP selection criteria, we generated two population structures and investigated their suitability for representing the panel.

Using all markers with a minor allele frequency (MAF) larger than 0.05^9,22,23^, reduced the total number of SNPs from 110,555 to 36,631, leading to the identification of five subpopulation clusters (Supplemental Fig. 2A). A second population structure was generated using stricter parameters, requiring the markers be biallelic, MAF > 0.05, one per gene and at least 500 kb apart. A total of 664 SNPs met these requirements, resulting in the identification of four subpopulation clusters (Supplemental Fig. 2B).

**Figure 2:**
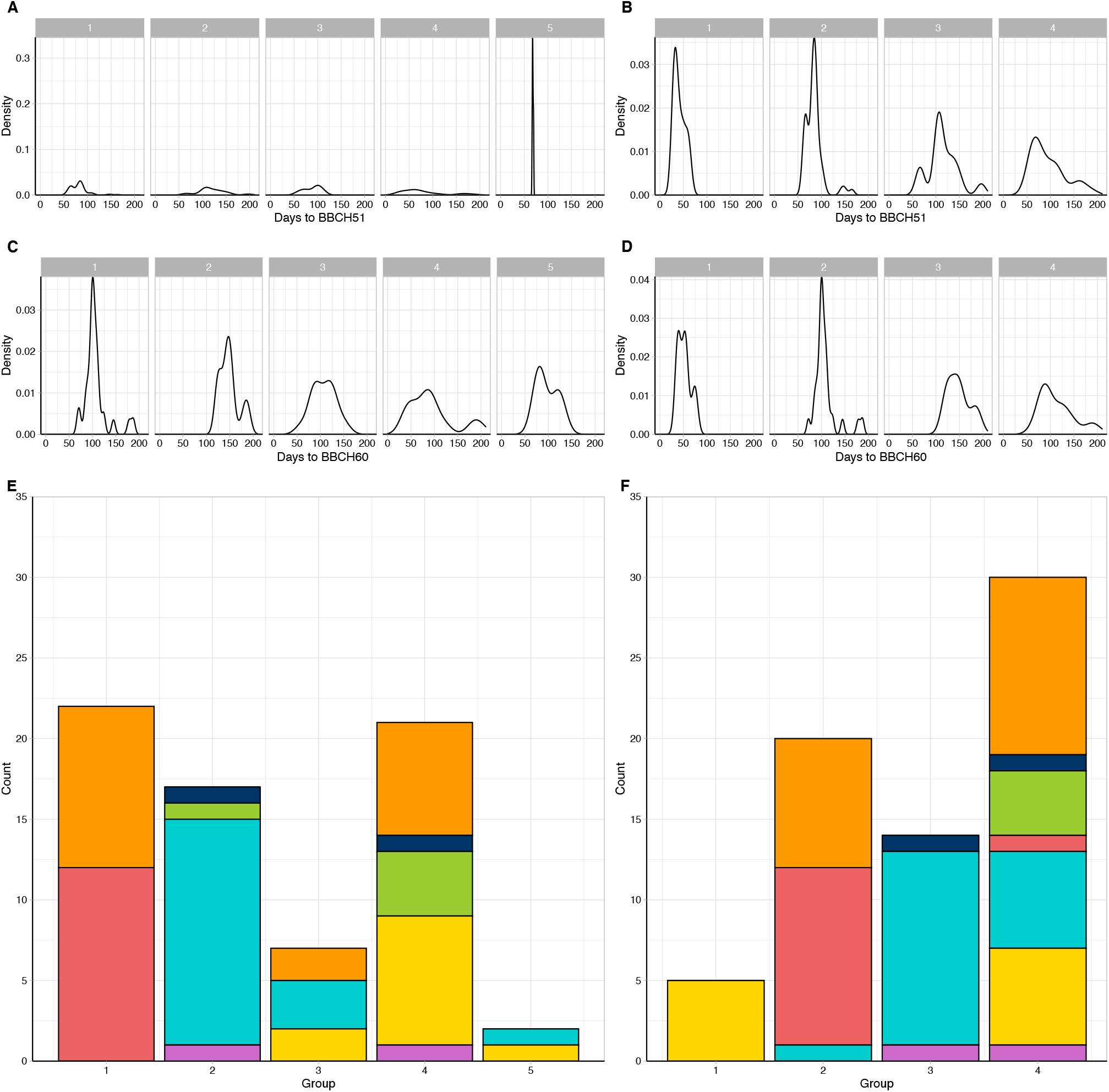
The choice of SNP pruning rules can significantly change the inferred population structure. Density plots representing A) DTB, C) DTF for the accessions within the five subpopulation clusters. Density plots representing B) DTB, D) DTF for the accessions within the four subpopulation clusters. E) Population structure generated from SNPs with MAF > 0.05 F) Population structure generated from more stringent SNP pruning (Biallelic only, MAF > 0.05, one per gene).

We assessed the two population structures based on crop type and phenotypic data. In the first population structure (generated using less stringent parameters), Fig. 2E, cluster one contained only broccoli and calabrese, both members of the same subspecies var. *italica*^24,25^, whereas cluster two was mainly cauliflower, subspecies var. *botrytis*, but including some late flowering accessions in both clusters. Interestingly, this population structure grouped the rapid cycling and late flowering kales together with a spread of accessions from other crop types, cluster four. The remaining two clusters were small by comparison: cluster three was comprised of seven accessions, a mixture of broccoli, cauliflower and kale; cluster five consisted of just two lines, one kale and one cauliflower.

The four clusters identified using more stringent SNP selection criteria contained all of the rapid cycling kales in cluster one, characterised by their early heading and flowering phenotypes (Figs. 2B, 2D, 2F). Cluster two was mainly broccoli and calabrese, whilst cluster three consisted largely of the earlier flowering cauliflowers. Cluster four contained the late flowering individuals from all crop types within the population, hence the larger variation in heading and flowering for this cluster.

Comparison of the clustering of accessions between the two population structures demonstrated the more stringent SNP criteria gave rise to a population structure in which individuals were grouped with other accessions that would be expected to be genetically similar based on knowledge of crop type and flowering phenotype. Consequently, this population structure was applied in subsequent GWAS analyses.

To gauge the extent of linkage disequilibrium we calculated the mean pairwise squared allele-frequency correlation (*r*^*2*^) for mapped markers. A linkage disequilibrium window of 50 (providing > 3 million pairwise values of *r*^*2*^) resulted in a mean pairwise *r*^*2*^ of 0.0979, confirming a low overall level of linkage disequilibrium in *B. oleracea*.

### Associative transcriptomics identifies miR172D as a candidate for controlling vernalisation response

SNP associations were compared to the physical positions of orthologues of genes known to be involved in the floral transition in Arabidopsis. A total of 43 flowering time related traits were analysed using this pipeline, including DTB and DTF for each treatment. A total of 111 significant SNPs were identified, P < 0.05, six of which demonstrated clear association peaks and were investigated further (Table 1).

**Table 1:** Significant SNP associations with vernalisation response in diverse *B. oleracea* accessions, detected across the genome (FDR < 0.05).

| Marker Information |  |  | Association Information |  |  |  |  |
| --- | --- | --- | --- | --- | --- | --- | --- |
| Marker | Chromosome | Alleles | -Log10(p) | Marker R<sup>2</sup> | Traits | Arabidopsis ID | Orthologue |
| Bo6g103650.1:2010:T | C06 | C/T/Y | 6.4017787 | 0.39231 | 6P 12V 10 °C BBCH51 | AT1G67140.3 | *SWEETIE* |
| Bo9g179000.1:2589:G | C09 | G/T/K | 6.4077566 | 0.39662 | 6P 12V 10 °C BBCH51 | AT5G04240.1 | *ELF6* |
| Bo1g011280.1:786:A | C01 | A/T/W | 6.0844894 | 0.44220 | 10P 12V 5 °C BBCH60 | AT4G31490.1 | Coatomer, beta subunit |
| Bo7g026810.1:124:G | C07 | A/G/R | 4.7781947 | 0.36476 | 6P 12V 10 °C BBCH60 | AT2G05790.1 | O-Glycosyl hydrolases family 17 protein |
| Bo7g104810.1:204:T | C07 | A/T/W | 5.9788107 | 0.41678 | 10P 6V 15-5 °C BBCH51 | AT3G55512 | mir172D |
| Bo2g009460.1:894:T | C02 | C/T | 7.6880767 | 0.40565 | 10P 6V 5 °C BBCH60-BBCH51 | AT5G10140.4 | *FLC.C2* |

We first sought to identify genetic associations with the trait data for the non-vernalized experiment. Whilst no significant association peaks were identified for DTB, an association at Bo8g089990.1:453:T was identified (P = 2.29E-06) for DTF under non-vernalizing conditions. This marker was within a region demonstrating good synteny to Arabidopsis, despite there being a number of unannotated gene models present. Conservation between Arabidopsis and *B. oleracea* suggests that this region contains an orthologue of microRNA172D, AT3G55512, which has been linked to the floral transition in *A. thaliana*^26,27^ (Fig. 3A). Furthermore, the difference in DTB between six and twelve weeks of vernalisation at 15 °C, identified a significant association on C07 at Bo7g104810.1:204:T (FDR, P < 0.05). This association was in the vicinity of a second orthologue of miR172D (Fig. 3C).

**Figure 3:**
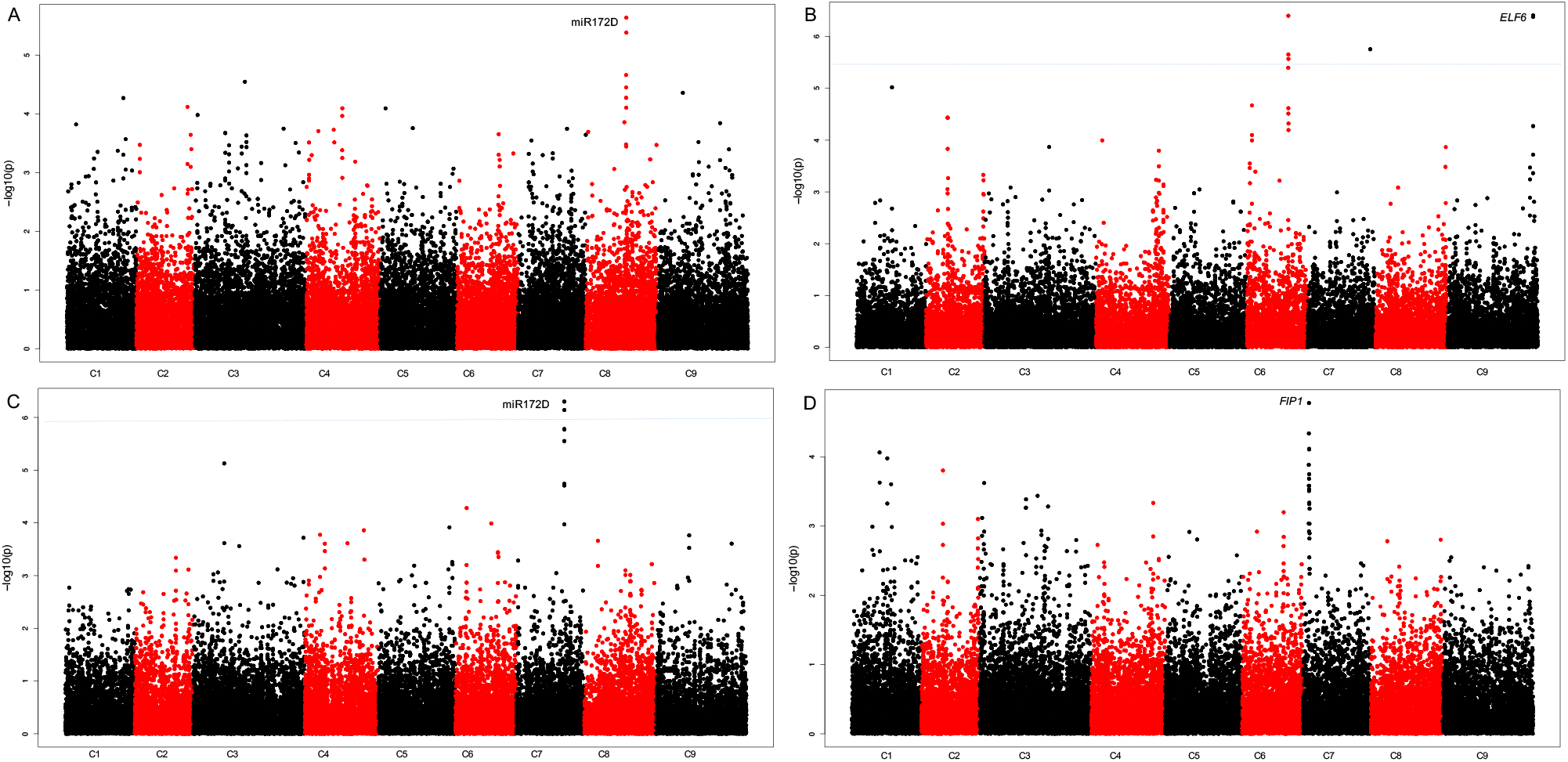
The developed pipeline identifies associations with flowering traits. Distribution of mapped markers associating with A) Number of DTF under non-vernalizing conditions B) DTB after a six-week pre-growth, twelve weeks vernalisation 10 °C C) The difference in DTB between six and twelve weeks of vernalisation at 15 °C, after exposure to a ten-week pre-growth D) The DTF after exposure to six-week pre-growth, twelve weeks vernalisation 10 °C. Sixty-nine accessions of *B. oleracea* were phenotyped for DTB and DTF and marker associations were calculated using a generalized linear model, implemented in TASSEL to incorporate population structure. Log_10_ (P values) were plotted against the nine *B. oleracea* chromosomes in SNP order. Blue line FDR threshold, P< 0.05.

We then analysed the association with traits relating to the timing of vernalisation. No significant associations were identified for traits after the six-week growth period followed by 5 °C vernalisation. However, a strong association was identified on C07 at the marker Bo7g026810.1:124:G, for DTF for the six-week growth period with twelve-weeks of vernalisation at 10 °C. Synteny with Arabidopsis suggests that an orthologue of *FRI INTERACTING PROTEIN 1*, (*FIP1*), AT2G06005.1 (Fig. 3D) is present within this region. Within Arabidopsis it has been demonstrated that *FIP1* interacts with *FRIGIDA* (*FRI*)^28^ which is a major source of natural variation in flowering time in Arabidopsis and has been shown to be important in determining vernalisation requirement. Additionally, significant associations (FDR, P < 0.05), were found for DTB for the six-week growth period with twelve weeks of vernalisation at 10 °C. An association was identified at Bo9g179000.1:2589:G, which is in the vicinity of an orthologue of *Early Flowering 6* (*ELF6*), AT5G04240.1 (Fig. 3B), a nuclear targeted protein able to affect flowering time irrespective of *FLC*.

The differences in flowering phenotype between the SNP variants for the four strongest associations were analysed (Fig. 4). There were significant differences in the traits associated with miR172D (DTF with no vernalisation and the difference in DTB for plants grown under vernalising temperatures of 5 and 15 °C) for different alleles (Fig. 4A and B). For the candidate identified through an association to the difference in DTB after exposure to vernalisation temperatures of 5 and 15 °C, five individuals contained the A variant, four of which were broccoli and one cauliflower. The alternate variant was a T allele and was present in 50 individuals. Conversely, the miR172D candidate identified through an association with DTF with no vernalisation, had 11 individuals with a C at this locus, whilst 51 had a T. Interestingly, the smaller set with the C allele still contained every crop type.

**Figure 4:**
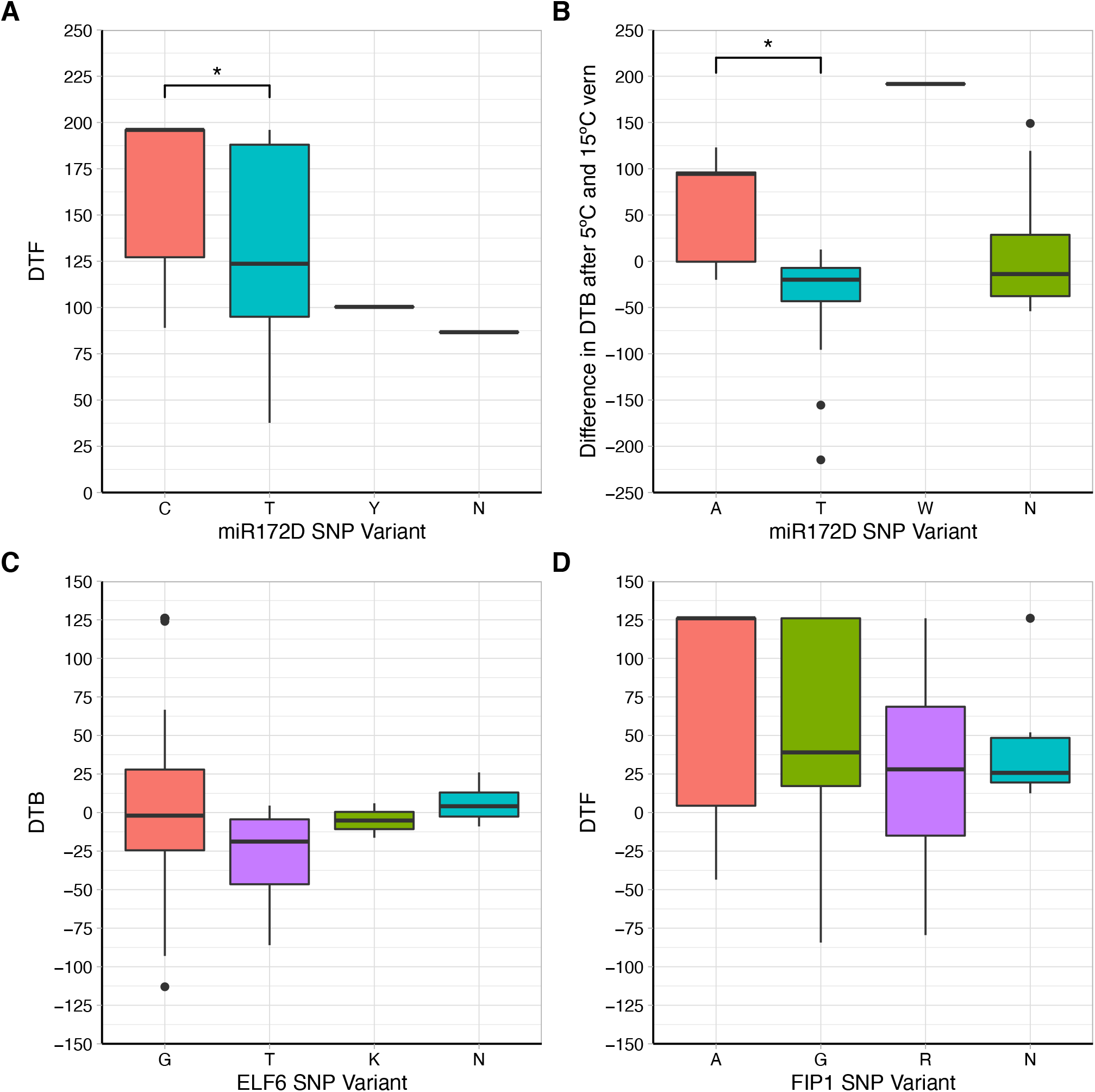
A significant phenotypic difference was found for individuals exhibiting SNP variants for the associations pointing to miR172D as a candidate. Boxplots represent the trait data, DTB or DTF for each of the significant markers alongside the different alleles present across the population for each marker. The box represents interquartile range, outliers are represented by black dots.

### GEM analysis identifies Bo*FLC*.*C2* as a candidate gene involved in the vegetative-to-floral transition

To investigate potential links between gene expression and the traits of interest, we performed gene expression marker (GEM) analysis. We identified Bo*FLC*.*C2* expression as being significantly associated with both the DTB and DTF under non-vernalising conditions (Fig. 5). Bo*FLC*.*C2* exhibited both low and high expression within the population. As expected, all five rapid cycling accessions demonstrated no Bo*FLC*.*C2* expression. Recently, a Brassica consortium developed targeted sequence capture for a set of relevant genes, including *FLC*. DNA from four of the five rapid cycling accessions had been enriched with that capture library and sequenced. Lacking a reference sequence for *B. oleracea* that contains *BoFLC*.*C2*, we used *B. napus* (cv. Darmor)^29^ as a reference to map the captured sequence data from the four rapid cycling accessions to. Comparison of *B. oleracea* transcript data^30^ to this Darmor genome reference revealed a 99.54 % identity in coding sequence, allowing Darmor to be used as a surrogate reference. Indeed, we found that all four rapid cycling accessions, GT050381, GT080767, GT100067 and GT110222, do not have *FLC*.*C2*, which was revealed by a lack of read mapping (Supplemental Fig. 5). Bo*FLC*.*C2* is known to be involved in vernalisation response^30^ and rapid cycling varieties do not require a period of vernalisation in order to transition to the floral state. As a control, we investigated mapping for 49 non-rapid cycling accessions where we expect *FLC*.*C2* to be present. For all 49 we found the expected read mapping evidence, confirming that use of the polyploid *B. napus* reference is appropriate (Suppl Fig. 7). We wouldn’t expect any one gene to explain all variation over the entire dataset for a complex trait and indeed only a weak positive correlation (DTB R^2^ = 0.024, DTF R^2^ = 0.036) between flowering phenotype and Bo*FLC*.*C2* expression was identified. However, a strong positive correlation (DTB R^2^ = 0.871, DTF R^2^ = 0.891) was found for the phenotypic extremes (the rapid cyclers with no expression and the later flowering lines with high levels of Bo*FLC*.*C2*), Fig.6, confirming a role for Bo*FLC*.*C2*.

**Figure 5:**
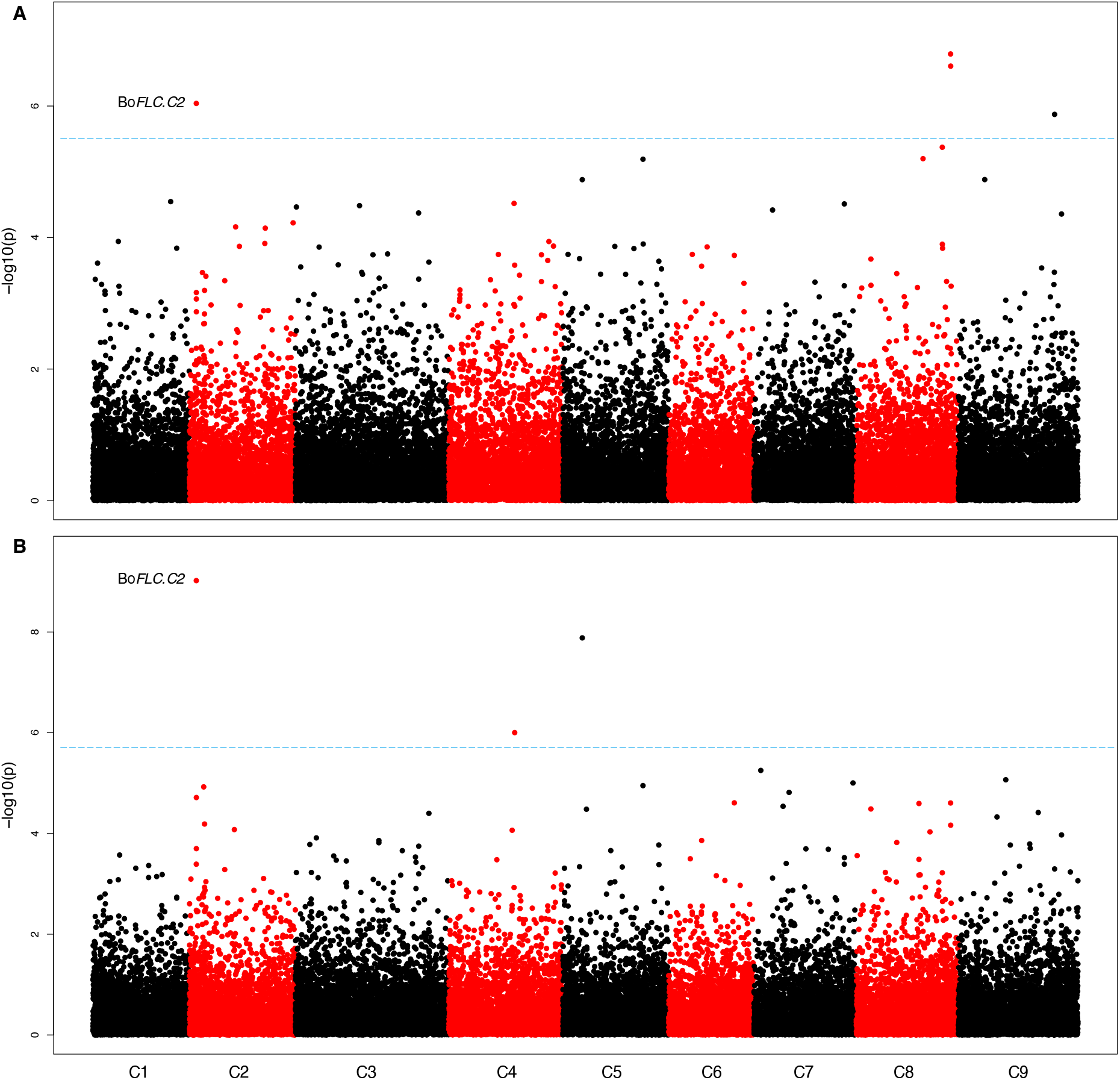
GEM analysis identifies *FLC* expression on chromosome C2 as a candidate for flowering traits under non-vernalising conditions. Distribution of gene expression markers associating with A) DTB after exposure to non-vernalizing conditions B) DTF after exposure to non-vernalising conditions. Log_10_ (P values) were plotted against the nine *B. oleracea* chromosomes in SNP order. Blue line FDR threshold, P < 0.05.

## Discussion

Producing synchronous *B. oleracea* vegetables is a key goal for growers and breeders. Quantifying vernalisation responses for different varieties is an important step towards this goal, providing a foundation for targeted breeding. Phenotyping for both DTB and DTF under different environmental conditions revealed a varied response within the population and identified some general trends. Altering the timing of vernalisation demonstrated that a shorter growth period prior to the exposure to cold extended the time taken to reach DTB and DTF.

This could be attributed to the presence of a juvenile phase in many of the lines, which has been widely documented in *B. oleracea*^16,31,32^. A juvenile plant is described as being unable to respond to floral inductive cues. The fact that many lines were able to flower much faster following longer pre-vernalisation growth, suggests they had reached the adult vegetative phase and were receptive to cold as a floral inductive cue. Further experimental work would be needed to test this hypothesis.

Increasing vernalisation length resulted, on average, in faster and more synchronous heading and flowering. This was a predicted outcome, as current knowledge suggests that increased vernalisation duration would saturate the vernalisation requirement of a larger proportion of the accessions. Furthermore, we found a positive correlation between vernalisation temperature and variation in heading and flowering. The 5 °C vernalisation period presented the most synchronous heading and flowering across all accessions, consistent with cooler temperatures meeting the vernalisation requirement for a higher proportion of the accessions.

In order to determine gene candidates underlying the observed differences in responses to vernalisation, we developed an associative transcriptomics pipeline for *B. oleracea*. To reduce the risk of false positives, we developed stringent criteria to identify unlinked markers for the determination of the population structure. The population structure was validated using crop type and phenotypic information on heading and flowering.

Using this validated population structure with associative mapping, we identified candidates orthologous to known Arabidopsis floral regulators, including miR172D. In Arabidopsis, the miR172 family post-transcriptionally supress a number of *APETALA1-*like genes, including *TARGET OF EAT1, 2* and *3*, which in turn aids the promotion of floral induction^27,33–35^. Furthermore, the SNP variant data for both of the associations that point to miR172D, exhibiting significant phenotypic differences. Two orthologues of Arabidopsis miR172D have been identified in *B. oleracea*^36^ but their functional roles have yet to be determined.

GEM analysis identified Bo*FLC*.*C2* expression as being significantly associated with both DTB and DTF under non-vernalising conditions. This can be attributed to the extreme phenotypes within the population (Fig. 6). No Bo*FLC*.*C2* expression was detected in five lines. A loss-of-function mutation at Bo*FLC*.*C2* in cauliflower has been associated with an early flowering phenotype^37^, indicating that Bo*FLC*.*C2* has an equivalent role in cauliflower to *FLC* in Arabidopsis. Four of the five lines for which Bo*FLC*.*C2* expression could not be detected did not have the Bo*FLC*.*C2* paralogue according to the EcoTILLING data. These four lines were all kales and demonstrated an early flowering phenotype, suggesting that Bo*FLC*.*C2* has a similar role to *AtFLC* also in kales, and potentially across *B. oleracea*. Although DTB and DTF were highly correlated with Bo*FLC*.*C2* expression under non-vernalising conditions for the phenotypic extremes, for the whole population the correlation was low. This is to be expected as Bo*FLC*.*C2* is just one of many genes that we expect to be involved in the floral transition within *B. oleracea* and therefore is unlikely to account for all the observed variation.

**Figure 6:**
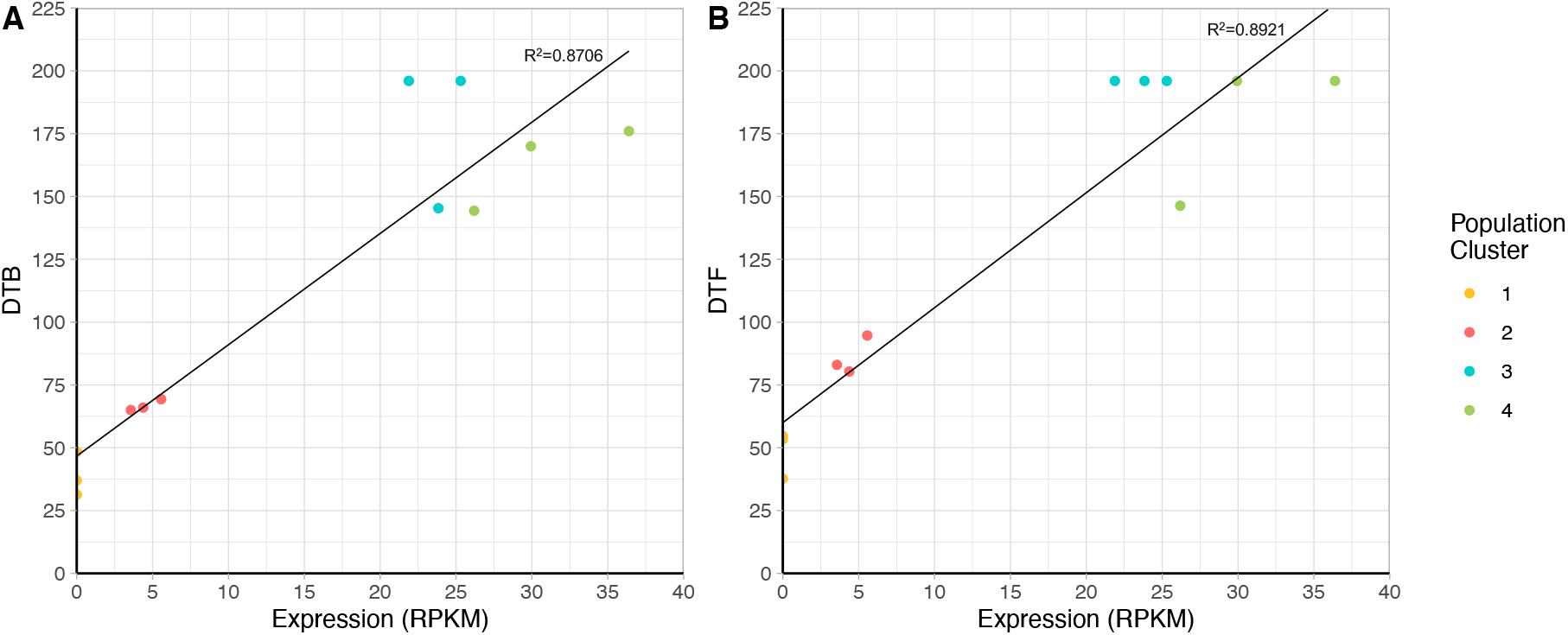
A strong positive correlation can be seen between lines at the phenotypic extremes and their Bo*FLC*.*C2* expression levels. Colours represent the subpopulation of each line, as determined by population structure analysis.

The expression data used for the GEM analysis was generated from leaf tissue at one timepoint. As a consequence, any genes which are not expressed in the leaf at this time will not be identified in this analysis. Use of transcriptome data from other tissues in addition to the leaf data could identify a greater number of associations.

## Conclusion

Identifying determinants of flowering time in *B. oleracea* is an important step in the production of synchronous brassica vegetables. Here, we generate and validate a novel pipeline for associative transcriptomics analysis in *B. oleracea* and show that this pipeline is effective in identifying genetic regulators of complex traits, such as flowering time. GWAS analysis identified miR172D as a candidate for vernalisation response, whilst GEM analysis identified a significant marker at *BoFLC*.*C2*, an important gene in the vernalisation pathway of *B. oleracea*. Our results provide insight into the genetic control of flowering in *B. oleracea*, and candidates which could provide a foundation for future breeding strategies.

## Methods

### Plant Materials and Growth Conditions

A subset of 69 lines fixed as doubled haploids (DH) or at S4 and above were chosen from the *Brassica oleracea* Diversity Fixed Foundation Set^15^. This subset contained accessions from seven different *B. oleracea* crop types. Seed was sown and seedlings pricked out into individual pots. Plants were watered daily, fed as required and given a pre-growth period of either six or ten weeks in a glasshouse. Natural light was supplemented with LED lights giving 16 h light (00:00 – 16:00) and 8 h dark. The daytime temperature was maintained at a minimum of 21 °C day, 18 °C night. At the end of the pre-growth period, three plants of each line for each of the pre-growth treatments were transferred to Conviron controlled environment rooms for a period of vernalisation at 5, 10 or 15 °C, 16 h LED light (00:00 – 16:00) and 8 h dark, 60% humidity. Vernalisation duration was either six or twelve weeks, after which plants were re-potted into 2 L pots and placed into a polytunnel using a randomised block design. Due to the staggered sowing for each experiment, all plants came out of vernalisation and into the polytunnel on the same day. Within the polytunnel, plants received natural light only. Additionally, three replicates of each line were grown without vernalisation as a non-vernalised control group. The plants were scored at buds visible (DTB) and upon opening of first flower (DTF)^38^.

### SNP Calling

The growth conditions, sampling of plant material, RNA extraction and transcriptome sequencing was carried out as described by He et al.^39^. The RNA-seq data from each accession were mapped on to CDS models from the *Brassica oleracea* pangenome^40^ as reference sequences, using Maq v0.7.1^41^. SNPs were called by the meta-analysis of alignments as described in Bancroft et al^42^. SNP positions were excluded if they did not have a read depth in excess of 10, a base call quality above Q20, missing data below 0.25, and 3 alleles or fewer.

### Population Structure and GWAS analyses

A population structure was generated from the 69 *B. oleracea* accessions used for phenotyping. Stringent rules were used to first prune the SNP data to ensure it only contained unlinked markers. SNPs were required to be biallelic, with a minor allele frequency (MAF) > 0.05 and a minimum distance of 500-bp between markers. The Bayesian clustering algorithms implemented in the software STRUCTURE^43^ were then used to determine population structure with a burn-in of 10000, a MCMC of 10000 and 10 iterations. STRUCTURE HARVESTER^44^ was used to determine the optimal *K* value, by generating a series of Δ*K* values, which represent the mean likelihood of *K* divided by the standard deviation of *K*, for the population. STRUCTURE HARVESTER was also used to produce the Q matrix used in GWAS analysis.

TASSEL^45^ version 5.0 was used to select the most appropriate model for each trait based on QQ plots. Generalised linear models (GLM), with correction for population structure using the Q matrix and PCA, and mixed linear models (MLM) with kinship data calculated with TASSEL’s ‘centered IBS’ method was used to look for associations. Optimum compression level and P3D variance component estimation were used as MLM options. For GWAS analysis only SNP markers with an allele frequency > 0.05 were used. To gauge the extent of linkage disequilibrium, the mean pairwise *r*^*2*^ was calculated using the SlidingWindow function within TASSEL version 5.0, with a linkage disequilibrium window of 50. SNPs were removed from GWAS analysis if their minor allele frequency was below 0.05.

To build a statistical model of the trait as a function of RPKM, gene expression marker (GEM) associations were calculated by an in-house script in R Version 3.6.3 using a fixed effect linear model with RPKM values. The script removed all markers with an average expression below 0.5 RPKM and performed a linear regression using RPKM as a predictor value to predict a quantitative outcome of the trait value. Both SNP and GEM outputs were plotted as Manhattan Plots created using an in-house R script.

Significance for both GWAS and GEM analyses was determined using false discovery rate (FDR). FDR is calculated on a trait by trait basis and uses the P-values from the GWAS or GEM analysis to determine the proportion false discoveries across all discoveries in the experiment^46^. FDR was calculated using the QValue package^47^ in R.

### DNA Extraction

Genomic DNA of accessions used in EcoTILLING was prepared from young leaf tissue of plants grown in a glasshouse. Plants were watered daily, fed as required and natural light was supplemented with LED lights giving 16 h light (00:00 – 16:00) and 8 h dark. The daytime temperature was maintained at a minimum of 21 °C day and 18 °C at night. Light was excluded for 48 h prior to harvesting. Approximately 3 g of tissue was used per accession and nuclei extraction buffer (10 mM Tris-HCL pH 9.5, 10mM EDTA pH 8.0, 100 mM KCL, 500 mM sucrose, 4 mM spermidine, 1 mM spermine, 0.1 % 2-mercaptoethanol) was added. The solution was vortexed and filtered through two layers of Miracloth. Lysis buffer (10 mM Tris-HCL pH 9.5, 10mM EDTA pH 8.0, 100 mM KCL, 500 mM sucrose, 4 mM spermidine, 1 mM spermine, 0.1 % 2-mercaptoethanol, 10 % Triton-x) was added to the homogenate, before centrifuging at 1000 g for 20 min at 4 °C. Supernatant was removed and the pellet resuspended in CTAB extraction buffer (100 mM Tris-HCL pH 7.5, 0.7 M NaCl, 10 mM EDTA pH 8.0, 1 % cetyl trimethylammonium bromide, 1 % 2-mercaptoethanol). The solution was incubated at 60 °C for 30 min. An equal volume of chloroform:isoamyl alcohol (24:1) was added and the solution rotated for 10 min before centrifugation for 10 min at 1000 g. The aqueous phase was removed and RNase T1 (50 units/ml) and RNaseA (50 μg/ml) added. The solution was incubated at 37 °C for 45 min. Proteinase K (150 μ/ml) was added and the solution incubated at 37 °C for a further 45 min. An equal volume of phenol:chloroform:isoamyl alcohol (25:24:1) and the solution centrifuged for 10 min at 1000 g, this was repeated once more. An equal volume of chloroform:isoamyl alcohol was added and the solution rotated for 10 mins, before centrifuging at 1000 g for 10 min. DNA was then precipitated by the addition of sodium acetate (10 % 3M) and 3 x volume of 100 % ethanol. The solution was centrifuged at 1000 g for 1 min to gently pellet the DNA and it was subsequently washed with 3 ml of 75 % ethanol. The solution was gently centrifuged, the ethanol poured off and the pellet left to air dry, before being resuspended in 50 μl dH_2_O. DNA was checked for quality. Nanodrop analysis was carried out on 1.5 μl of each DNA sample and the extracted DNA was stored at −20°C until required.

### Targeted Sequence Enrichment analysis

A bait library for targeted sequence enrichment for a specific subset of genes was developed and synthesized with Arbor Biosciences (https://arborbiosci.com/). DNA (Supplemental Table 1) was extracted and sent to Arbor for enrichment and Illumina sequencing. Reads from individual accessions were mapped to the reference sequence of *B. napus* cv. Darmor-*bzh*^29^ using BWA^48^ version 0.7.17-r1188. For BWA, we used aln/sampe and standard parameters. Mapped reads were sorted and indexed using SAMTOOLS^49^ version 1.10, sort and index and subsequently visualized with Integrative Genomics Viewer (IGV)^50^.

## Supplemental information

**Supplemental Table 1:** List of accessions with associated crop type information that were used for EcoTILLING analysis.

**Supplemental Table 2**: Analysis of the smaller SNP data set with the Bayesian clustering algorithms implemented in the program STRUCTURE identified four population clusters.

**Supplemental Figure 1:** Increased synchrony in DTB and DTF was observed as vernalisation temperature was reduced. Histograms representing the distribution of DTB and DTF post-vernalisation across the population after exposure to vernalization at 5, 10 or 15 °C. Individuals that did not flower have been removed from this plot.

**Supplemental Figure 2**: *ΔK* based on rate of change of LnP, Maxima indicates the DeltaK that best explains the population structure. Plots produced using STRUCTURE Harvester output. A) DeltaK values for biallelic SNPs, MAF > 0.05, one SNP per gene, >500kb apart, *K* = 4. B) DeltaK values calculated for SNPs with MAF > 0.05, *K* = 5.

**Supplemental Figure 3**: Quantile-Quantile Plots for SNP associations with A) the DTF under NV conditions. GLM, with Q matrix correction for population structure B) the DTB after six-week pre-growth and 10°C vernalisation for twelve-weeks. GLM, with Q matrix correction for population structure C) The difference in DTB following 5 °C and 15 °C vernalisation for six-weeks, after exposure to a ten-week pre-growth. GLM with Q matrix correction for population structure D) The DTF after exposure to six-week pre-growth, twelve weeks vernalisation 10 °C. GLM with PCA correction for population structure.

**Supplemental Figure 4:** Linkage disequilibrium decay. A) Bo8g089990.1:453:T, miR172D candidate. B) Bo9g179000.1:2589:G, *ELF6* candidate. C) Bo7g026810.1:124:G, *FIP1* candidate. D) Bo7g104810.1:204:T, miR172D candidate.

**Supplemental Figure 5:** Mapping Bo*FLC*.*C2* using Darmor-*bzh* as a reference. Four rapid cycling accessions and three representative accessions for the rest of the population.

## Acknowledgements

We thank Profs Lars Ostergaard and Steve Penfield (JIC) and Drs Andrea Harper and Lenka Havlickova (York) for discussion and critical comments on the manuscript. We thank Dr Alex Calderwood (JIC) for guidance and advice. We also thank University of Warwick for use of the *B. oleracea* Diversity Fixed Foundation Set. SW was supported by the UK Biotechnology and Biological Sciences Research Council (BBSRC) NRPDTP PhD Studentship Programme. JI and RW acknowledge funding from BBSRC Institute Strategic Programme (BB/P013511/1), RM, JI and RW acknowledge support from the BBSRC strategic LoLa ‘Brassica Rapeseed and Vegetable Optimisation’ (BB/P003095/1) and IB and ZH acknowledge funding from BBSRC (BB/L002124/1). RM acknowledges support from EU-Horizon2020 ERC Synergy Grant ‘PLAMORF’ (ID: 810131). Additional funding was provided by BBSRC sLoLa ‘Renewable Industrial Products from Rapeseed’ (BB/L002124/1).

## Author contributions

JI, RM, RW and SW designed the experiments that were carried out by SW with support from JI and RW. The SNP calling was carried out by ZH under guidance of IB. SW performed the phenotyping of material, all analyses and produced all figures. RW, IB and WH provided genomics and bioinformatics advice. HW provided programming support and guidance. BS designed and constructed the bait library for targeted sequence enrichment and carried out subsequent sequence mapping, which was analysed by SW. SW drafted the manuscript which was planned and refined by SW, RW and All authors contributed to writing the manuscript.

