## Supplemental Table 1 for "Validation of a novel associative transcriptomics pipeline in *Brassica oleracea:* Identifying candidates for vernalisation response"

| Genotype | Crop Type |
|---|---|
| DJ3290 | Brussels Sprouts |
| GT050381 | Kale |
| GT060871 | Broccoli |
| GT070808 | Cabbage |
| GT071233 | Broccoli |
| GT080340 | Broccoli |
| GT080638 | Calabrese |
| GT080713 | Cauliflower |
| GT080760 | Kale |
| GT080767 | Kale |
| GT080843 | Calabrese |
| GT080869 | Cauliflower |
| GT080876 | Cauliflower |
| GT080891 | Kale |
| GT081012 | Kale |
| GT081137 | Cauliflower |
| GT081140 | Broccoli |
| GT081150 | Broccoli |
| GT081395 | Broccoli |
| GT090058 | Cabbage |
| GT090341 | Cauliflower |
| GT100062 | Broccoli |
| GT100065 | Broccoli |
| GT100067 | Kale |
| GT110222 | Kale |
| GT110231 | Cauliflower |
| GT110244 | Broccoli |
| GT110251 | Cauliflower |
| GT110257 | Calabrese |
| GT110266 | KohlRabi |
| GT120144 | Broccoli |
| GT120152 | Cauliflower |
| GT120155 | Kale |
| GT120160 | Calabrese |
| GT120163 | Calabrese |
| GT120168 | Brussels Sprouts |
| GT120170 | Cauliflower |
| GT120182 | Cauliflower |
| GT120192 | Cauliflower |
| GT120194 | Calabrese |
| GT120201 | Calabrese |
| GT120205 | KohlRabi |
| GT120208 | Broccoli |
| GT120210 | Cabbage |
| GT120211 | Calabrese |
| GT120213 | Cauliflower |
| GT120214 | Broccoli |
| GT120215 | Broccoli |
| GT120218 | Cauliflower |
| GT120223 | Cauliflower |
| GT120226 | Cabbage |
| GT120231 | Calabrese |
| GT120233 | Broccoli |
