## Supplemental Table 2 for "Validation of a novel associative transcriptomics pipeline in *Brassica oleracea:* Identifying candidates for vernalisation response"

| Genotype | CropType | Q1 | Q2 | Q3 | Q4 |
|---|---|---|---|---|---|
| DJ3290 | Brussels Sprouts | 0.002 | 0.002 | 0.002 | 0.995 |
| GT081137 | Cauliflower | 0.01 | 0.005 | 0.001 | 0.984 |
| GT050381 | Kale | 0.995 | 0.001 | 0.001 | 0.003 |
| GT060867 | Kale | 0.934 | 0.002 | 0.002 | 0.062 |
| GT060871 | Broccoli | 0.015 | 0.003 | 0.002 | 0.979 |
| GT110231 | Cauliflower | 0.001 | 0.001 | 0.997 | 0.001 |
| GT070808 | Cabbage | 0.035 | 0.002 | 0.012 | 0.952 |
| GT080340 | Broccoli | 0.01 | 0.005 | 0.006 | 0.979 |
| GT080486 | Kale | 0.022 | 0.002 | 0.005 | 0.97 |
| GT071233 | Broccoli | 0.001 | 0.899 | 0.098 | 0.002 |
| GT080713 | Cauliflower | 0.001 | 0.002 | 0.996 | 0.001 |
| GT080760 | Kale | 0.008 | 0.002 | 0.002 | 0.988 |
| GT080767 | Kale | 0.99 | 0.002 | 0.002 | 0.006 |
| GT080843 | Calabrese | 0.001 | 0.997 | 0.001 | 0.001 |
| GT080849 | Broccoli | 0.001 | 0.997 | 0.001 | 0.001 |
| GT080869 | Cauliflower | 0.019 | 0.008 | 0.374 | 0.598 |
| GT080876 | Cauliflower | 0.003 | 0.014 | 0.976 | 0.006 |
| GT080891 | Kale | 0.018 | 0.045 | 0.19 | 0.747 |
| GT081012 | Kale | 0.181 | 0.022 | 0.017 | 0.78 |
| GT081062 | Cauliflower | 0.004 | 0.003 | 0.907 | 0.087 |
| GT081103 | Broccoli | 0.003 | 0.005 | 0.003 | 0.989 |
| GT081140 | Broccoli | 0.002 | 0.981 | 0.011 | 0.005 |
| GT081150 | Broccoli | 0.001 | 0.997 | 0.001 | 0.001 |
| GT081395 | Broccoli | 0.021 | 0.043 | 0.019 | 0.916 |
| GT090058 | Cabbage | 0.003 | 0.002 | 0.001 | 0.994 |
| GT090341 | Cauliflower | 0.001 | 0.001 | 0.997 | 0.001 |
| GT100062 | Broccoli | 0.002 | 0.968 | 0.02 | 0.009 |
| GT100065 | Broccoli | 0.004 | 0.355 | 0.223 | 0.419 |
| GT100067 | Kale | 0.992 | 0.001 | 0.003 | 0.003 |
| GT080638 | Calabrese | 0.005 | 0.988 | 0.004 | 0.003 |
| GT110206 | Cauliflower | 0.001 | 0.002 | 0.992 | 0.004 |
| GT110222 | Kale | 0.992 | 0.002 | 0.004 | 0.002 |
| GT110244 | Broccoli | 0.009 | 0.843 | 0.142 | 0.006 |
| GT110251 | Cauliflower | 0.001 | 0.005 | 0.987 | 0.006 |
| GT110257 | Calabrese | 0.001 | 0.975 | 0.001 | 0.023 |
| GT110266 | KohlRabi | 0.002 | 0.003 | 0.992 | 0.003 |
| GT110275 | Kale | 0.002 | 0.009 | 0.002 | 0.986 |
| GT120144 | Broccoli | 0.013 | 0.139 | 0.161 | 0.687 |
| GT120152 | Cauliflower | 0.002 | 0.003 | 0.565 | 0.43 |
| GT120155 | Kale | 0.007 | 0.003 | 0.009 | 0.98 |
| GT120160 | Calabrese | 0.004 | 0.974 | 0.019 | 0.003 |
| GT120162 | Calabrese | 0.003 | 0.979 | 0.015 | 0.002 |
| GT120163 | Calabrese | 0.004 | 0.126 | 0.003 | 0.868 |
| GT120164 | Broccoli | 0.3 | 0.028 | 0.005 | 0.668 |
| GT120168 | Brussels Sprouts | 0.001 | 0.001 | 0.998 | 0.001 |
| GT120170 | Cauliflower | 0.018 | 0.011 | 0.882 | 0.089 |
| GT120179 | Cauliflower | 0.003 | 0.005 | 0.459 | 0.533 |
| GT120182 | Cauliflower | 0.002 | 0.088 | 0.33 | 0.58 |
| GT070654 | Cauliflower | 0.002 | 0.006 | 0.606 | 0.385 |
| GT120194 | Calabrese | 0.002 | 0.845 | 0.15 | 0.004 |
| GT120195 | Calabrese | 0.001 | 0.995 | 0.002 | 0.002 |
| GT120198 | Cauliflower | 0.02 | 0.004 | 0.473 | 0.504 |
| GT120201 | Calabrese | 0.002 | 0.991 | 0.004 | 0.002 |
| GT120205 | KohlRabi | 0.008 | 0.065 | 0.018 | 0.909 |
| GT120208 | Broccoli | 0.004 | 0.184 | 0.26 | 0.553 |
| GT120210 | Cabbage | 0.007 | 0.009 | 0.049 | 0.935 |
| GT120211 | Calabrese | 0.001 | 0.996 | 0.002 | 0.001 |
| GT120213 | Cauliflower | 0.001 | 0.001 | 0.997 | 0.001 |
| GT120214 | Broccoli | 0.004 | 0.649 | 0.012 | 0.335 |
| GT120215 | Broccoli | 0.002 | 0.629 | 0.005 | 0.363 |
| GT120218 | Cauliflower | 0.002 | 0.002 | 0.993 | 0.003 |
| GT120222 | Cauliflower | 0.004 | 0.696 | 0.213 | 0.087 |
| GT120223 | Cauliflower | 0.004 | 0.137 | 0.336 | 0.523 |
| GT120225 | Calabrese | 0.001 | 0.981 | 0.016 | 0.002 |
| GT120226 | Cabbage | 0.003 | 0.004 | 0.008 | 0.985 |
| GT120231 | Calabrese | 0.001 | 0.997 | 0.001 | 0.002 |
| GT120233 | Broccoli | 0.001 | 0.018 | 0.002 | 0.978 |
| GT120234 | Broccoli | 0.003 | 0.006 | 0.002 | 0.989 |
| GT120235 | Broccoli | 0.001 | 0.003 | 0.007 | 0.989 |
