## Supplementary figures and images for "Validation of a novel associative transcriptomics pipeline in *Brassica oleracea:* Identifying candidates for vernalisation response"

### Supplemental Figure S1

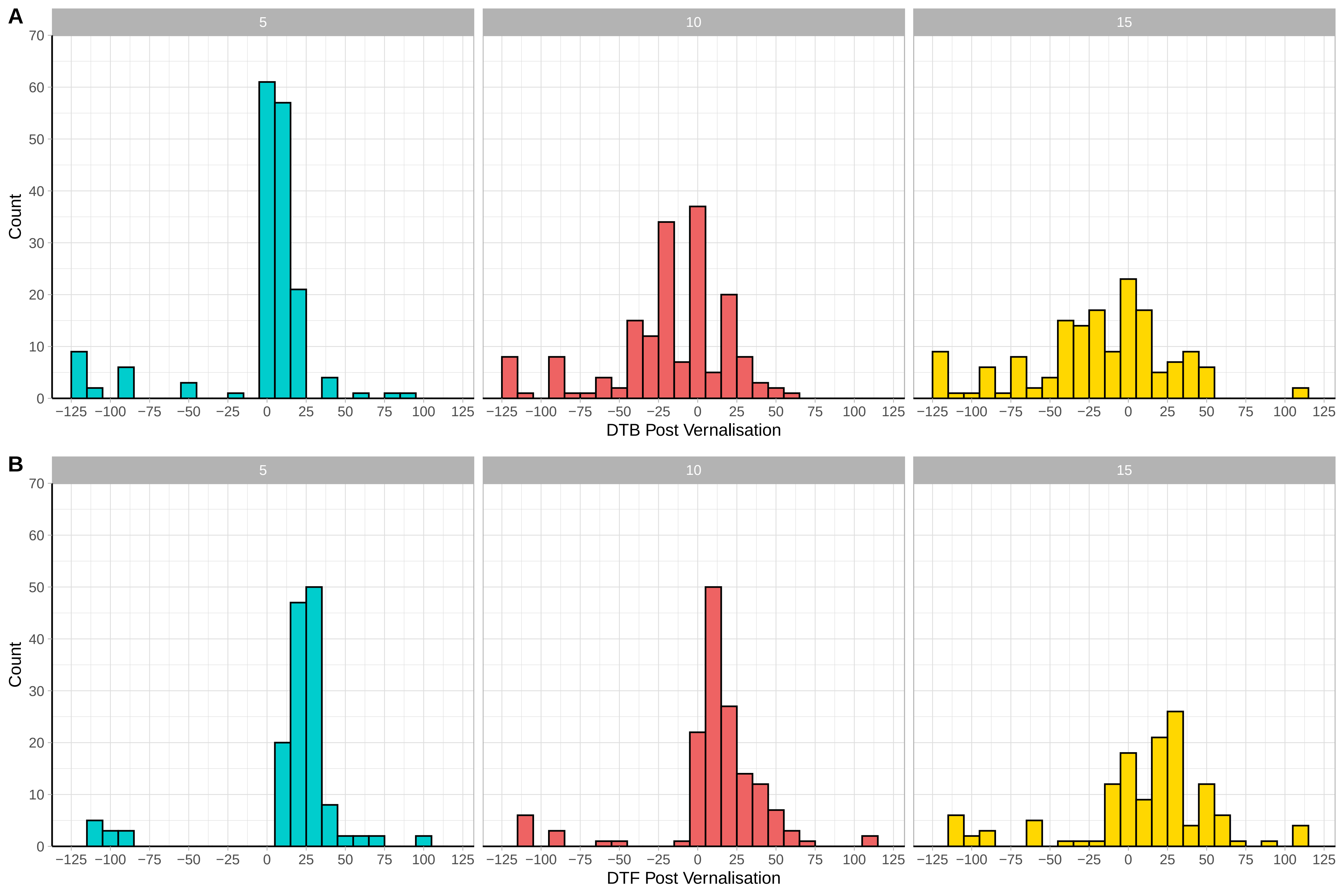

### Supplemental Figure S2

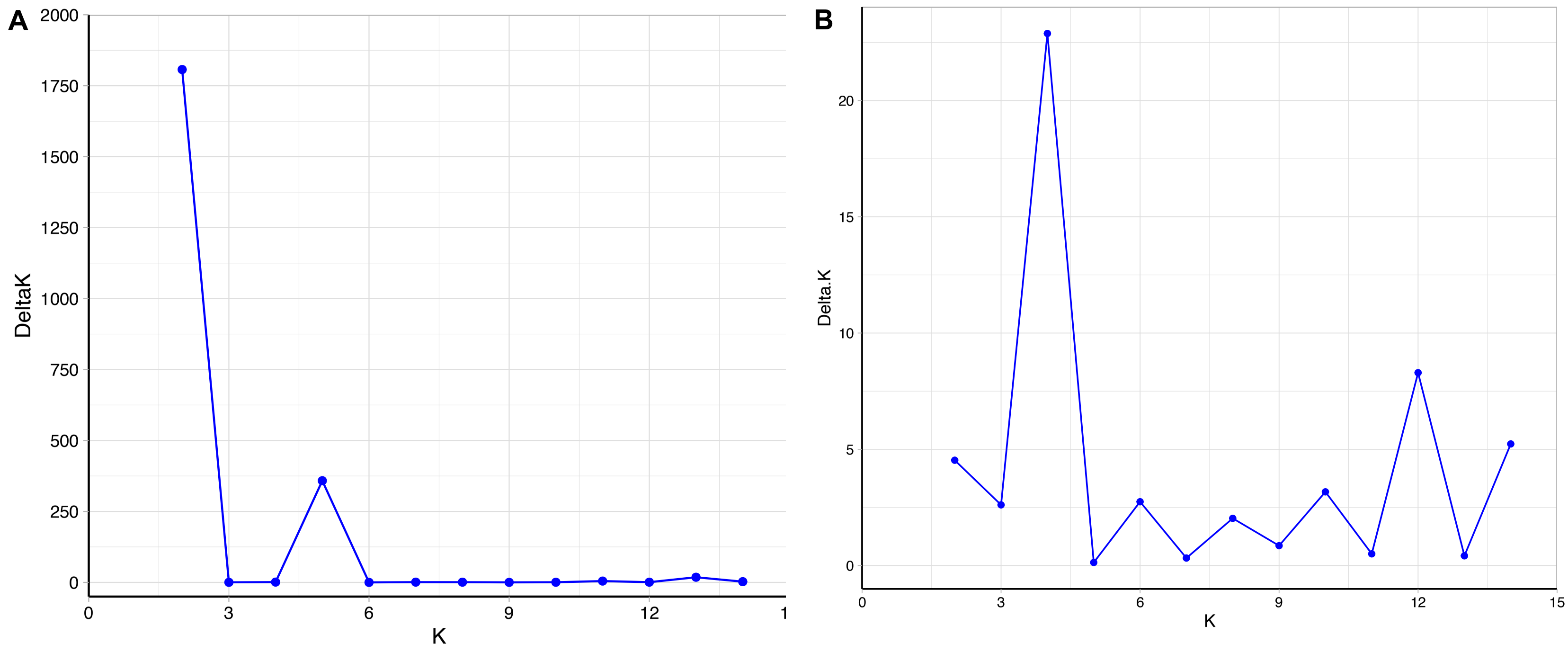

### Supplemental Figure S3

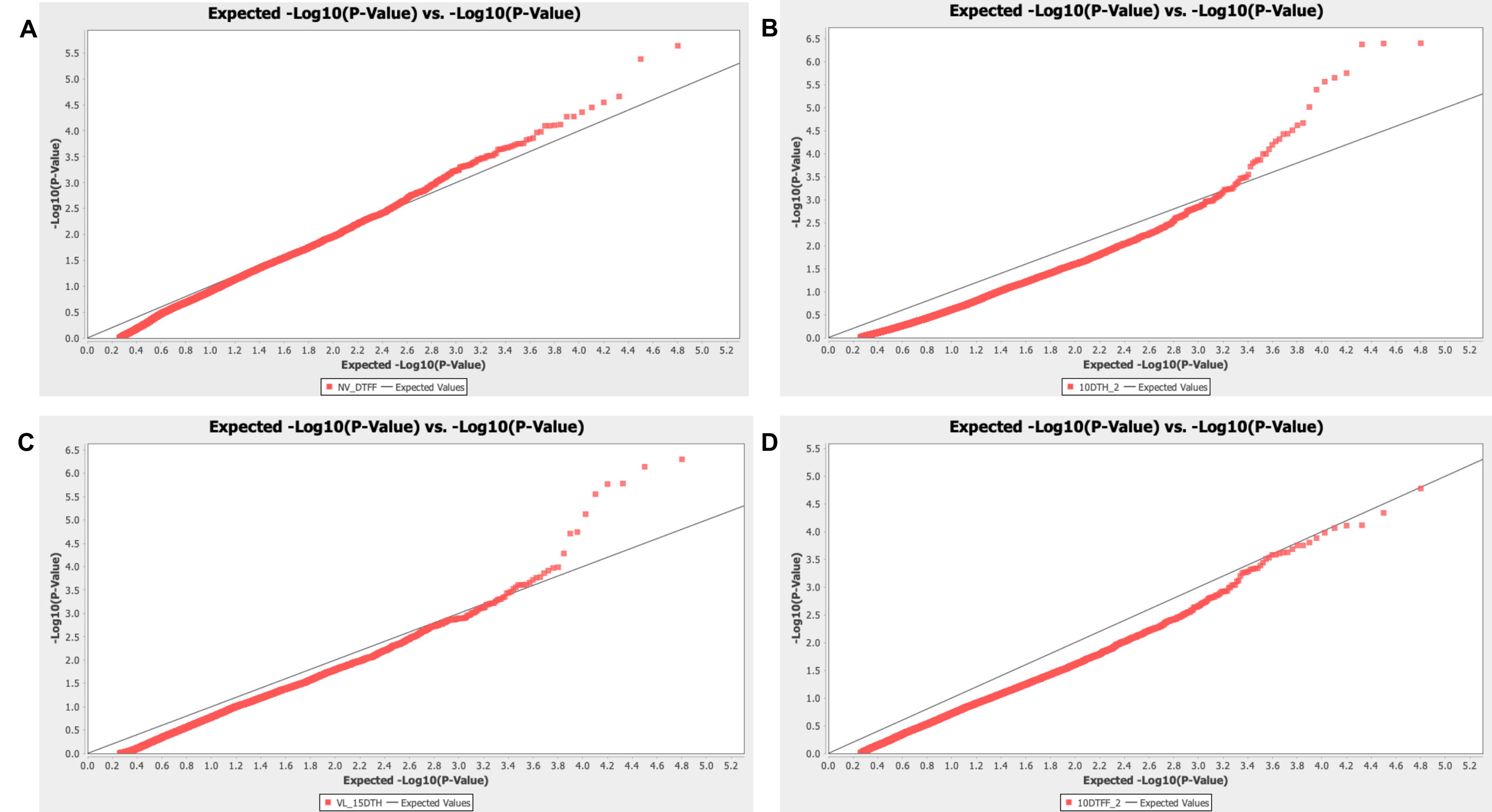

### Supplemental Figure S4

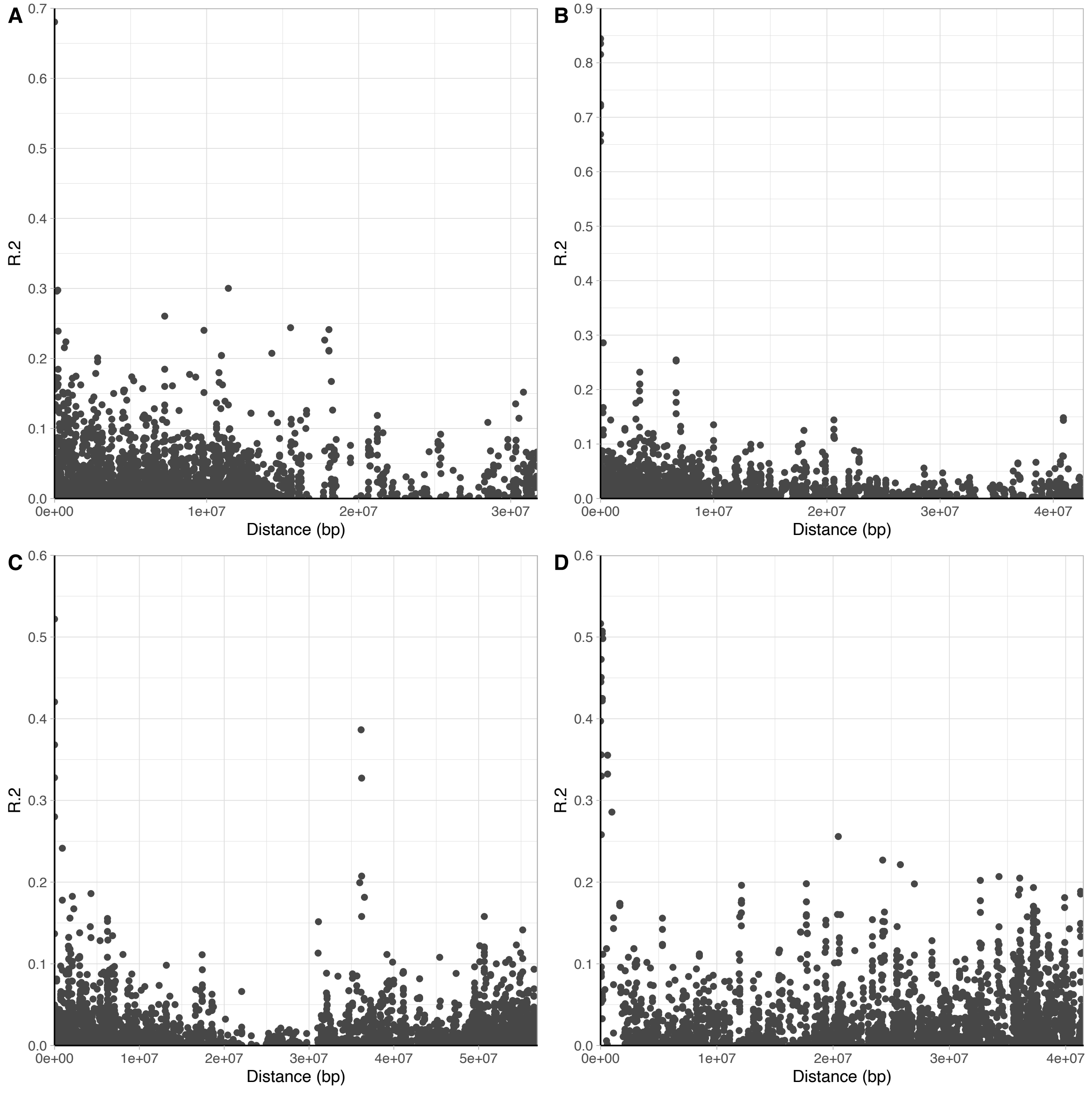

### Supplemental Figure S5

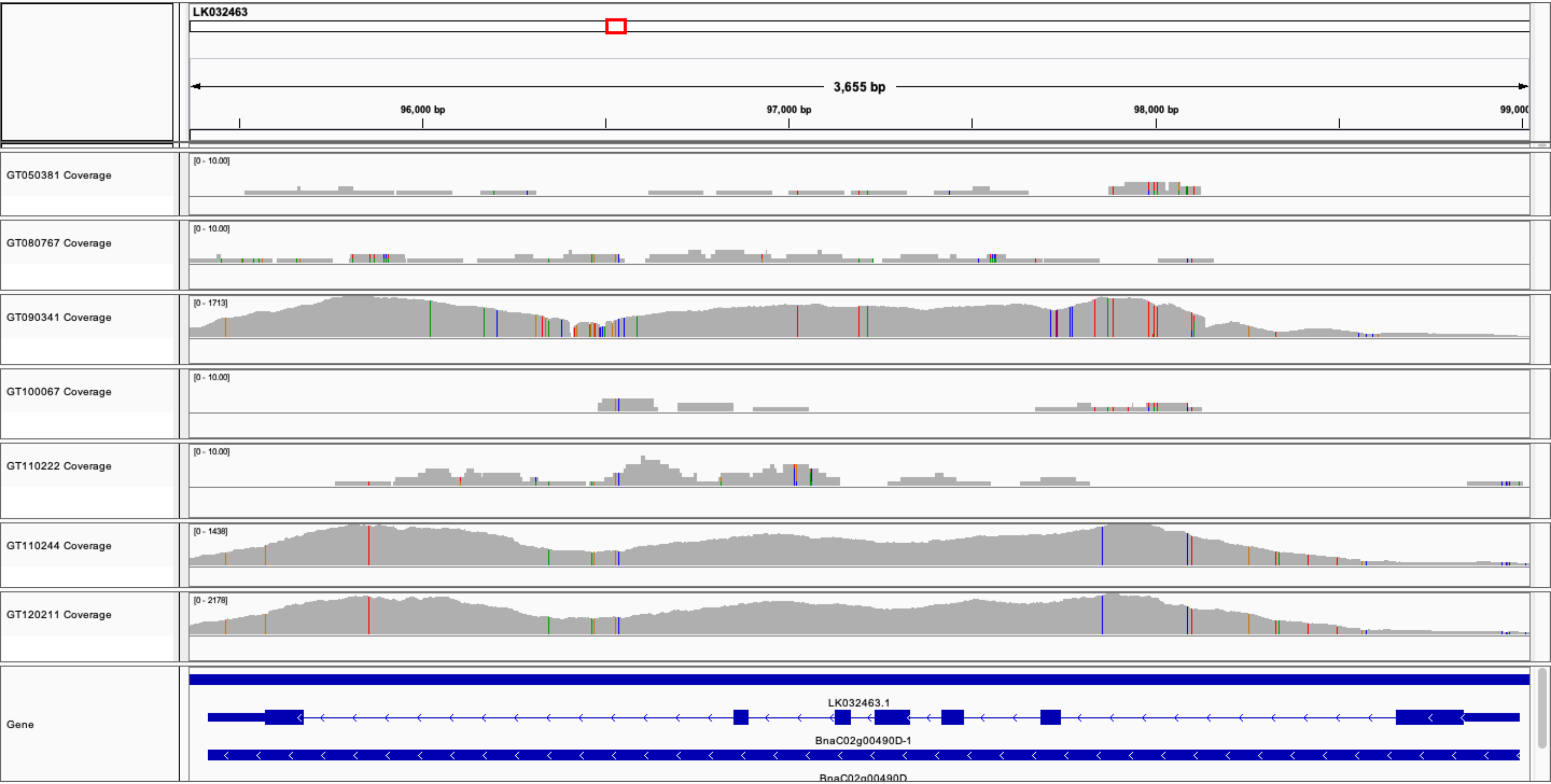
